# Microbiome transfer from native to invasive species may increase invasion risk and shorten invasion lag

**DOI:** 10.1101/2023.08.28.555072

**Authors:** Maria M. Martignoni, Oren Kolodny

## Abstract

In a fast-changing world, understanding how organisms adapt to their environment is a pressing necessity. Research has focused on genetic adaptation, while our understanding of non-genetic modes is still in its infancy. Particularly, the host-associated microbiome may strongly influence an organism’s ability to cope with its environment. The presence of certain microbes in the gut, for example, can facilitate the utilization of dietary resources, provide protection from pathogens, and increase resilience to diverse abiotic conditions. However, the role that the microbiome may play in species’ adaptation to novel challenges is largely unexplored, experimentally as well as theoretically. Here, we study the possibility of such adaptation in invasive species. We present and explore a new hypothesis: Invasive species may rapidly adapt to local conditions by adopting beneficial microbes of similar co-occurring native species. Ironically, due to competition, these native species are also those most likely to suffer from the invaders’ spread. We formulate a mathematical framework to investigate how the transfer of beneficial microbes between a native and an introduced species can alter their competitive dynamics. We suggest that, non-intuitively, the presence of a related native species may *facilitate* the success of an invasive species’ establishment. This occurs when the invader’s fitness is strongly influenced by adaptation to local conditions that is provided by microbes acquired from the natives’ microbiomes. Further, we show that in such cases a delayed acquisition of native microbes may explain the occurrence of an invasion lag, and we discuss biological systems that could lend themselves for the testing of our hypotheses. Overall, our results contribute to broadening the conceptualization of rapid adaptation via microbiome transfer and offer possible insights for designing early intervention strategies for invasive species management during their lag phase.

## Introduction

Invasive species cause annual damages of billions of dollars (Haubrock et al., 2021; Pimentel et al., 2005; Paini et al., 2016), and understanding the factors facilitating their adaptation is paramount for mitigating their impact. Early detection and eradication of potential invaders has been regarded as the cheapest and most effective control strategy (Epanchin-Niell, 2017), where an interesting phenomenon in particular may offer opportunities for early intervention: the occurrence of invasion-related lags (Crooks, 2005; Simberloff, 2003). An *invasion lag* is a prolonged period of time which is sometimes observed between the establishment of an alien species and the time point at which it becomes invasive, rapidly increasing in numbers and spreading geographically. This phenomenon has been documented for a large number of invasive plants (Aikio et al., 2010; Leung et al., 2012), invertebrates (Yanygina, 2017), birds (Aagaard and Lockwood, 2014), fishes (Azzurro et al., 2016), amphibians (Toledo and Measey, 2018), and reptiles (Guerrero et al., 2013), with invasion lag times lasting for years or even decades in some cases. To date, the underpinnings of invasion lags are little understood, and accordingly they are not predictable, rendering innocuous species and species that will become invasive indistinguishable (Coutts et al., 2018).

Several theories have been proposed to explain the occurrence of invasion lags (Simberloff, 2013). For instance, changes in the biotic or abiotic environment can affect the invasion dynamics (Crooks, 2005). Thus, a herbivore might keep an alien species under control, and its removal might allow it to rapidly spread unchecked (Strauss, 2014). Changes in climate might also affect invaders’ activity and community structure (Stachowicz et al., 2002; Wallingford et al., 2020), and human activity might at some point create conditions which are more favorable for invasion, allowing a seemingly-benign established alien species to suddenly become invasive (Fausch et al., 2001; Lee et al., 2021).

A perhaps more intriguing type of dynamics that can determine the length of an invasion lag, facilitating a switch from a low-frequency alien species with a limited spread to an invasive species with significant impact on the ecosystem, may stem from changes in the invasive population itself. One such possibility is via introduction of a new variant of the established species, which is coincidentally better adapted to local conditions or adds to the founder population the genetic diversity necessary to overcoming inbreeding depression (Dlugosch and Parker, 2008; Kolbe et al., 2004; Frankham, 2005). However, we now have evidence that variation can also emerge within the founder population itself, which becomes more successful over time as it evolves in the new environment (Prentis et al., 2008). Thus, genetic or phenotypic adaptation may provide the necessary fitness advantage to the introduced species, increasing its invasion success (Whitney and Gabler, 2008). This phenomenon has mostly been documented in plants (Matesanz et al., 2010; Ayres et al., 2004; Colautti et al., 2009), but it has also been observed in animals (Colautti and Lau, 2015), e.g. cane toads in Australia have evolved increasingly longer legs, accelerating their invasive spread (Phillips et al., 2006), and phenotypic plasticity has been found to contribute to invasion success in social insects (Manfredini et al., 2013, 2019).

Here, we propose an alternative explanation for the occurrence of invasion-related lags. Namely, we consider the possibility that adaptation in invasive species can be conferred by the acquisition of beneficial microbes. Beneficial microbes may in principle be acquired from the new environment that the invasive species reach. We suggest that this is unlikely, because environmental microbes would rarely be able to survive within a healthy host, and even if they do, these facultative associations are likely to be of secondary importance for fitness compared to co-evolved relationships.

Instead, we suggest that a likely source of beneficial microbes are native hosts. We analyze the case in which invaders become better adapted to local conditions through the acquisition of mutualistic microbes from the microbiome of phylogenetically and ecologically similar cooccurring native species. Phylogenetic closeness increases the likelihood that microbes that may have co-evolved locally with native hosts are pre-adapted to establishing a similar mutualistic relationships with introduced hosts. Ecological similarities reflect in similar basic needs of the native and invasive species, and thus in native microbes having a similar adaptive potential for invasive species.

It is increasingly recognized that host-microbiome interactions can shape host fitness and evolutionary potential, e.g., by increasing host tolerance to abiotic stress, by allowing the breakdown of local food sources, or by protecting the host from pathogens (Kolodny et al., 2020; Fontaine et al., 2022; Kikuchi et al., 2012; Kohl et al., 2014; Townsend et al., 2019; Fontaine and Kohl, 2020; Chiu et al., 2017; Kolodny and Schulenburg, 2020). Importantly, this response can be extremely rapid. For example, a reduction in microbiome diversity in the gut of tadpoles can decrease host fitness and its tolerance to thermal stress within days (Fontaine et al., 2022), and the acquisition of a pesticide degrading bacteria can confer on bean bugs immediate resistance to pesticides (Kikuchi et al., 2012). We also know that microbiome transmission is rarely strictly vertical, but can occur horizontally from a host to another through different pathways, such as direct contact between individuals, coprophagy (i.e. eating other individual’s feces), predation of younger individuals, or pickup of microbes that survive an intermediate phase in the environment outside the host (Robinson et al., 2019; Kolodny et al., 2019). Thus, the horizontal transfer of beneficial microbes from natives to invaders may facilitate their rapid adaptation, providing them with a fitness advantage and increasing their competitive ability.

Although the field of microbial ecology is growing rapidly, the current literature has focused on understanding how the presence or absence of certain microbes may affect host fitness (Fontaine et al., 2022; Kikuchi et al., 2012; Kohl et al., 2014; Fontaine and Kohl, 2020), without explicitly considering the ecological consequences of this fitness advantage. Only few studies have explored the influence of microbiome-related dynamics on multi-species communities (Martignoni et al., 2023, 2020; Daybog and Kolodny, 2022), where studies considering how variations in microbial communities may affect invasion have primarily dealt with the transmission of pathogens, rather than with the exchange of mutualistic microbes (Gruber et al., 2019; Faillace et al., 2017). Here we present a theoretical framework to investigate the possibility that microbiome sharing, between and within species, would alter the dynamics of invasion by conferring rapid ecological adaptation to invaders. In particular, we analyse how different characteristics of the native and invasive populations, such as their growth rate, carrying capacity and competitiveness, interplay with the probability of acquiring beneficial microbes to determine invasion success. We will discuss the role of microbiome transfer in determining lag times in biological invasion, and we will provide concrete directives on how our hypotheses may be tested in simple experimental settings.

In this study we focus on the transmission of beneficial microbes, however if invasive hosts can acquire beneficial microbes from natives, we hould expect that native hosts would also be able to acquire microbes from invasive hosts. Additionally, the transferred microbes may not be necessarily beneficial, and could be neutral or harmful to their new host (Dickie et al., 2017; Bahrndorff et al., 2016; Henry et al., 2021; Goss et al., 2020). Full treatment of these dynamics is beyond the scope of the current paper and are treated in a separate manuscript (Martignoni et al., in preparation).

## Model and Methods

We formulate an ordinary differential equation model to study the coupled dynamics of a native population *N* competing with an introduced population *I*, whereby interactions are modelled according to the competitive Lotka-Volterra equations (Gilad, 2008). The populations experience logistic growth until reaching a certain carrying capacity, where competition between species can reduce or even reverse the growth (see supplementary information, section A for a complete mathematical analysis of the Lotka-Volterra equations).

We consider that a beneficial microbiome can be transferred from native to introduced individuals and we explore the system’s dynamics under a range of parameters that govern this process. We consider that horizontal transmission can occur directly, through contact among individuals, or indirectly, with transmission mediated by the environment. This may include, for example, cases in which individuals of the invasive species utilize roosts or shelters that were previously occupied by native hosts, cases of coprophagy, or situations where birds of the different species share sites of sand or water bathing. We also posit that, once acquired, the microbiome may be vertically and horizontally transferred within the introduced population. In our study we do not differentiate between microbes, nor between their locations within the host. Rather with ‘microbiome’ we mean any collection of symbiotic microorganisms that increases fitness in its host. For simplicity we treat the transmission of the microbiome as a single event which may or may not occur, although in reality we expect transmission of only few microbial species - but with potentially large effects on fitness.

We model this scenario of interest by splitting the introduced population *I* into two sub-groups: the subpopulation that has not acquired microbes from native hosts (*I*_0_) and the subpopulation that has acquired microbes from native hosts (*I*_*m*_). Individuals can move from *I*_0_ to *I*_*m*_ by acquiring native microbes through interaction with natives (*N*), or through interaction with introduced individuals that have already acquired native microbes (*I*_*m*_). Sub-populations *I*_0_ and *I*_*m*_ compete for space, as the overall size of the introduced population is limited by a fixed carrying capacity. Mathematically, we write:

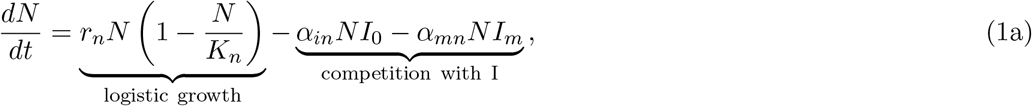

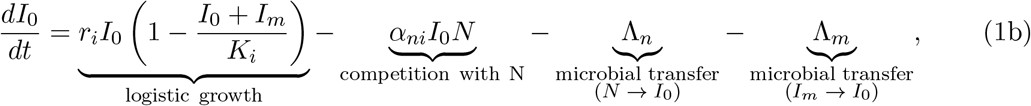

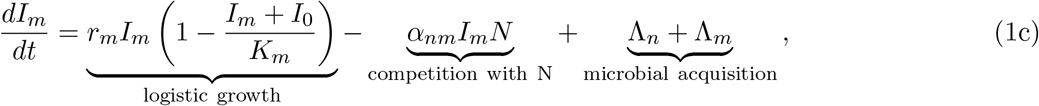

The ability of a population *j* to outcompete population *w* depends on its growth rate (*r*_*j*_), on its carrying capacity (*K*_*j*_), and on the competitive effect of population *w* on *j* (*α*_*wj*_). With ‘competitive ability’ we refer therefore to the set of traits of a population (in our model, the set of parameters *r*_*j*_, *K*_*j*_ and *α*_*wj*_) that characterize the growth of population *j* in the presence of population *w*, with populations *j* and *w* being the native and introduced populations. The population that outcompetes the other is referred to as the ‘superior competitor’.

If the waiting time for a microbiome transfer event to happen is exponential, as commonly assumed in modelling, microbiome transfer can be simulated as a Poisson process with a rate which depends on the density-dependent microbiome transfer rate from natives to introduced individuals (*λ*_*n*_), and on the size of the native and introduced populations (*N* (*t*) and *I*_0_(*t*) respectively). This implies that the number of introduced individuals that acquire native microbes in the time interval (*t, t* + *dt*] through interspecific contact is a Poisson random variable Λ_*n*_(*t*), with rate *γ*_*n*_(*t*) = *λ*_*n*_*N* (*t*)*I*_0_(*t*), such that:

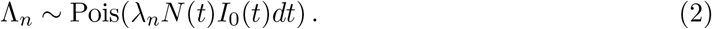

The density-dependent microbiome transfer rate *λ*_*n*_ may depend on the factors which underlie the biology of transmission and host-microbe interactions. For instance, ecological similarity or phylogenetic relatedness between native and invasive hosts may increase the likelihood that native microbes may establish in an invasive host (Rojas et al., 2021; Parker et al., 2015). Parameter *λ*_*n*_ may also depend on the mode of transmission: Direct contact between hosts, e.g., through predation, may increase the likelihood of microbial acquisition by a new host, while indirect contact, e.g. through the use of the same sand or water for bathing or digging, may lead to a lower rate of microbial acquisition. Finally, *λ*_*n*_ may depend on the microbes themselves, as not all microbes are equally likely to be transmitted or acquired (Moeller et al., 2018).

Once acquired, the microbiome can be transferred horizontally from *I*_*m*_ to *I*_0_ through the same modalities described above, at a rate which depends on the density-dependent microbiome transfer rate among introduced individuals (*λ*_*m*_) and on the size of subpopulations *I*_0_ and *I*_*m*_. Again, as for *λ*_*n*_ the value of parameter *λ*_*m*_ should also depend on the mode of transmission and on the characteristics of the transferred microbes. The number of individuals that acquire native microbes through intraspecific contact can be described by a Poisson random variable Λ_*m*_(*t*), with rate *γ*_*m*_(*t*) = *λ*_*m*_*I*(*t*)*I*_0_(*t*) such that

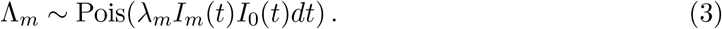

We will look at the situation in which a small number of introduced individuals are released into the environment while the native population is at its carrying capacity, and we will discuss scenarios in which the introduced population is poorly adapted to local conditions before acquiring native microbes, and better adapted after. Mutualistic microbes can increase host access to new food resources, increase host growth, or decrease its mortality, e.g., by protecting the host from pathogens (Qu et al., 2020; Raymann and Moran, 2018). We characterize therefore the subpopulation that has acquired native microbes (*I*_*m*_) by a higher carrying capacity and/or a higher growth rate with respect to the subpopulation that has not acquired native microbes (*I*_0_), and consider all the possible emerging dynamics. The mathematical analysis of scenarios A-C is presented in the supplementary information, sections B and C. Simulations are run in MATLAB2022b using the Euler’s Method, and the code is publicly available at https://github.com/nanomaria/microbiometransfer.

### Scenario A The timing of microbiome acquisition affects invasion lag times

Prior to microbiome transfer, the native and the introduced population have reached an equilibrium of stable coexistence (i.e., *K*_*n*_ *< r*_*i*_*/α*_*ni*_ and *K*_*i*_ *< r*_*n*_*/α*_*in*_), whereby introduced individuals are few with respect to natives (Fig. S2, scenario A). This scenario is realized in the model when the introduced population has a lower carrying capacity than the native population, but a higher growth rate or a strong competitive effect on natives (i.e., *r*_*i*_ *> r*_*n*_ or *α*_*in*_ *> α*_*ni*_). The acquisition of a native microbes leads to an increase in the carrying capacity of the introduced population, which becomes competitively superior and grows larger, displacing (or reducing the size of) the native population. If we consider that invaders and natives can coexist for a long time before microbiome is transferred from a species to the other, analysis of this scenario provides insights into the possible role of microbiome-mediated adaptation in driving a lag in biological invasion, and into the impact of horizontal microbiome transfer between and within species on the invasion lag time.

### Scenario B The establishment of an introduced species is made possible by transfer of microbes from native species

The introduced population cannot stably establish in the new environment and experiences a population decline after introduction, due to not being able to attain positive population growth (modelled as considering *r*_*i*_ *<* 0 and *dI*_0_*/dt* = *r*_*i*_*I*_0_, see supplementary information, section D), or due to competition with well-adapted natives (i.e., *K*_*n*_ *> r*_*i*_*/α*_*ni*_ and *K*_*i*_ *< r*_*n*_*/α*_*in*_). We consider that the adoption of native microbes leads to an increase in the growth rate and/or carrying capacity of the introduced population, rescuing it from extinction. Analysis of this scenario provides insights into the probability that an introduced population will adapt and stably establish in a new environment thank to the transfer of microbes from natives. Ironically, after having acquired native microbes, the now adapted introduced population increases in size, causing the native population to decline, or even go extinct (Fig. S2, scenario B).

### Scenario C: The presence of natives facilitates adaptation

Introduced individuals are superior competitors, and in the presence of an introduced population natives are driven to extinction (i.e., *K*_*i*_ *> r*_*n*_*/α*_*in*_ and *K*_*n*_ *< r*_*i*_*/α*_*ni*_). However, the introduced population also has a low carrying capacity (Fig. S2, scenario C), due to being poorly adapted to the local conditions. Adaptation can occur through the acquisition of native microbes, which in our simulations causes an increase in the carrying capacity of the introduced population. This means that if introduced individuals acquire the microbiome from natives before displacing them through competition, the introduced population will thrive, otherwise their population size will remain small. Analysis of this scenario provides insights into the interplay of competitive ability, patch size, and population densities in determining the circumstances under which microbiome-mediated adaptation is most likely to occur.

## Results

### Scenario A The timing of microbiome acquisition affects invasion lag times

If competition between natives and introduced individuals is weak, stable coexistence is observed, whereby the size of the introduced population is small due to being poorly adapted to local conditions. A situation of stable coexistence is maintained until the first microbiome transfer event between species occurs, and a new subpopulation *I*_*m*_ is created, which is better adapted and has a higher competitive ability than the introduced subpopulation *I*_0_ (Fig. 1a). In this simulation we consider vertical transmission of microbes to occur faithfully between parent and offspring, and we set horizontal transmission to zero. Thus, growth in the competitively superior *I*_*m*_ population through reproduction leads to the exclusion of natives. Given that the overall size of the invasive population is limited by a fix carrying capacity, the better adapted subpopulation *I*_*m*_ will eventually also displace subpopulation *I*_0_ through competition for space.

**Fig. 1:**
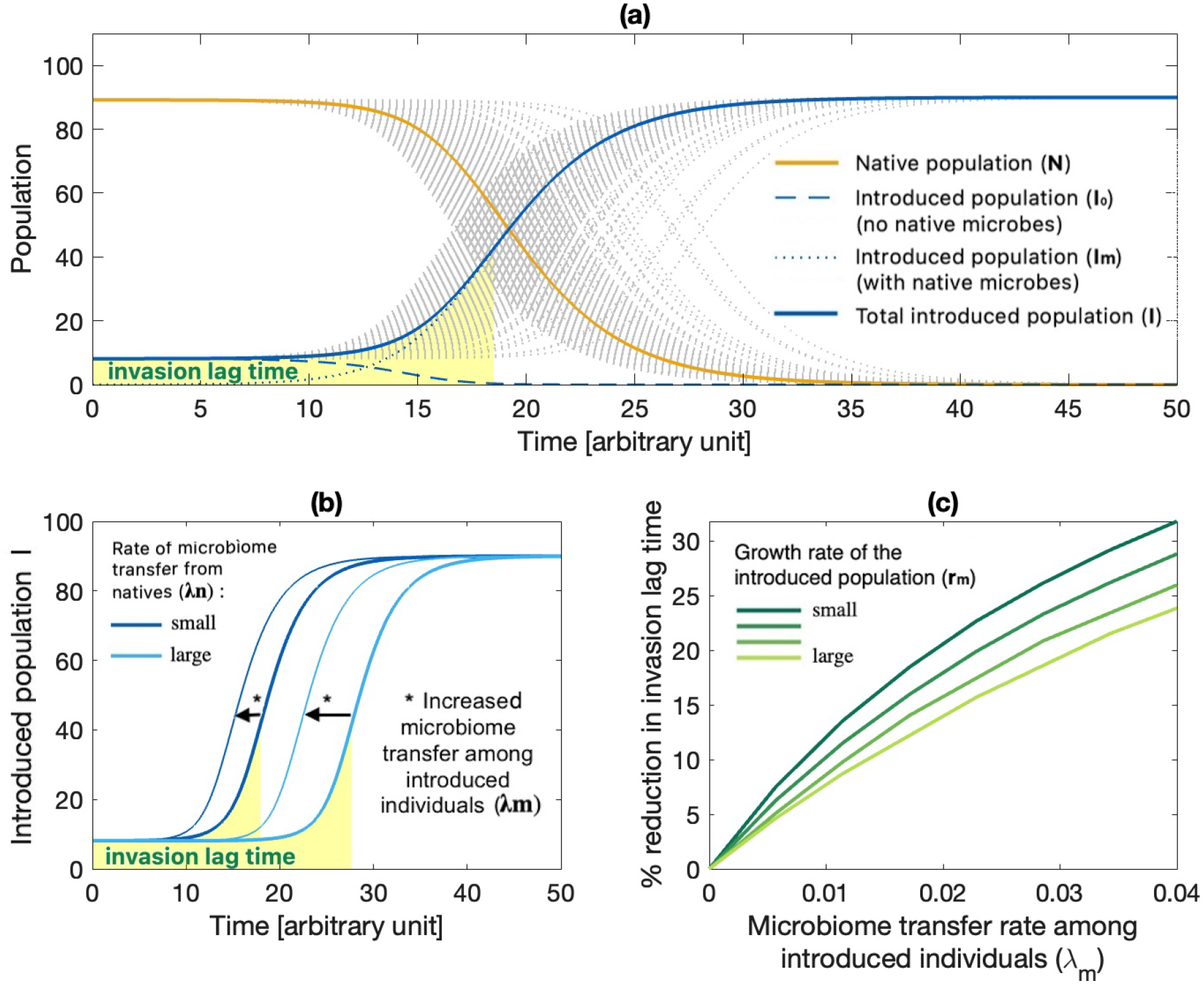
(a) A native population *N* (orange curve) coexists with a much smaller introduced population *I*_0_ (blue curve). Microbiome transfer from native to introduced individuals leads to adaptation of the introduced population (formulated in the model as the creation of a new subpopulation *I*_*m*_, where *I*_*m*_ + *I*_0_ represents the total introduced population *I*), which becomes competitively superior and invasive, and displaces the native population. The invasion lag time is shaded in yellow, and it is defined as the time interval from species introduction to when the population growth of the introduced species reaches its inflection point. Grey dotted lines correspond to the results of 500 stochastic realizations of Eq. (1), while the orange and blue solid curves correspond to their mean average. (b) The invasion lag time increases when microbiome transfer events from natives to introduced individuals are rare (i.e., when the density-dependent microbiome transfer rate *λ*_*n*_ is small, light blue curve) and decreases when events are more frequent (i.e., when *λ*_*n*_ is large, dark blue curve). The occurrence of horizontal microbiome transfer among introduced individuals (i.e., between the subpopulations *I*_0_ and *I*_*m*_, parameter *λ*_*m*_) can decrease the invasion lag time. (c) Percent reduction in invasion lag time as a function of the horizontal microbiome transfer rate among introduced individuals (*λ*_*m*_), for different growth rates of the subpopulation *I*_*m*_ (*r*_*m*_). The contribution of horizontal microbiome transfer in reducing the invasion lag time is larger when the growth rate *r*_*m*_ is small. Note that figures (b)-(c) consider the solutions corresponding to the mean average of a large number of stochastic realizations of Eq. (1).

We call ‘invasion lag time’ the time interval occurring between species introduction and the inflection point in the population growth of the introduced species, which depends on the time of the first microbiome transfer event. The lower the rate of microbiome transfer from natives to introduced individuals, the longer the invasion lag time, where horizontal microbiome transmission among introduced individuals can speed up the spread of beneficial microbes within a population, and decrease the invasion lag time (Fig. 1b). This effect is particularly prominent when the growth rate of the introduced population is low compared to the rate of horizontal transmission (Fig. 1c). In this case it will take longer for the subpopulation with native microbes (*I*_*m*_) to competitively displace the subpopulation without native microbes (*I*_0_). Thus, subpopulation *I*_0_ will still be largely represented in the total introduced population, slowing down the population growth of the introduced population as a whole, unless horizontal microbiome transfer among introduced individuals leads to a quick spread of native microbes, and to a direct conversion of *I*_0_ into *I*_*m*_.

### Scenario B The establishment of an introduced species is made possible by transfer of microbes from native species

If the introduced population is poorly adapted to the local conditions, it may experience a decline in its population size after introduction, due to its own inability to sustain a positive population growth or due to competition with better adapted natives. Microbiome transfer from natives can facilitate the adaptation of the introduced species (by increasing its growth rate and/or carrying capacity) and ease its establishment. Thus, under this scenario, the stably establishment of an introduced population is made possible by transfer of microbes from native species (Fig. 2a).

**Fig. 2:**
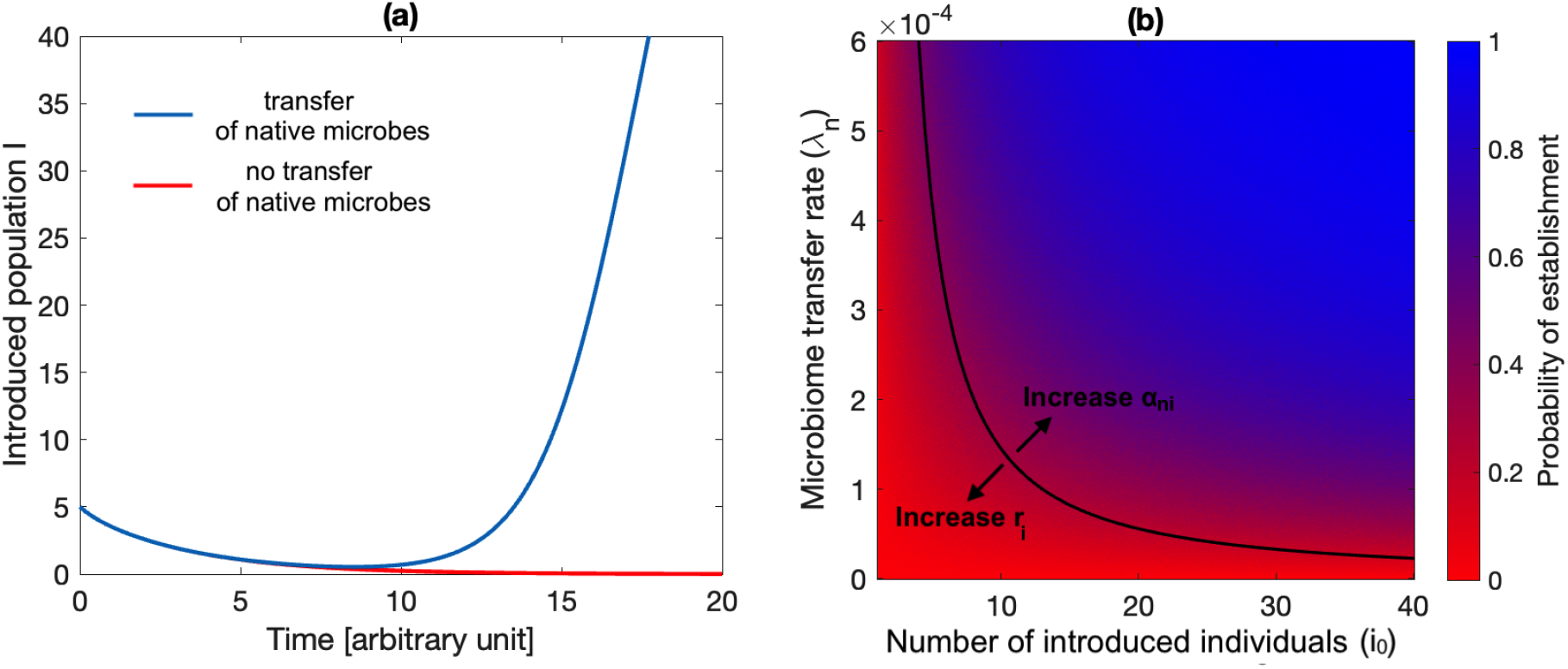
(a) A poorly adapted introduced population fails to establish if it does not timely acquire beneficial microbes from co-occurring natives (red curve). The transfer of beneficial microbes from natives to introduced individuals can confer adaptation to local conditions to the introduced population and rescue it from extinction (blue curve). (b) The probability of establishment of an introduced population (computed as the mean of 500 realizations) increases with increasing number of introduced individuals *i*_0_, and with increasing density-dependent rate of microbiome transfer between the native and introduced population *λ*_*n*_. The black curve represents the deterministic approximation derived in Eq. (4). Increasing the competitive effect of natives on introduced individuals *α*_*in*_ reduces their probability of establishment, while a larger growth rate of the introduced population *r*_*i*_ increases it.

Interestingly, the same native population that facilitates the establishment of an introduced species is subsequently likely to suffer from its spread. Indeed, after establishing the now adapted introduced population experiences a rapid population growth, which may coincide with a reduction in the population size of natives (Fig. S2, scenario B). If population growth occurs only after a long lag time, the resulting dynamics of invasion is similar to the lagged invasion discussed in scenario A (cfr. Fig. 1a and Fig. S4a).

Increasing the number of introduced individuals increases the probability that a timely microbiome transfer event will occur and facilitate the establishment of the introduced population (Fig. 2b). There are two reasons for this increase: (i) if the number of introduced individuals is large, it will take longer to the poorly adapted introduced population to die out, which increases the probability that a microbiome transfer event will occur in time to confer adaptation to local conditions to the introduced population before its extinction; and (ii) a large introduced population increases the rate of possible encounters between natives and introduced individuals, making a microbiome transfer event more likely to occur.

In the supplementary information (section D), we derive a condition to determine under which circumstances microbiome exchange can facilitate the establishment of a poorly adapted introduced population. We obtain:

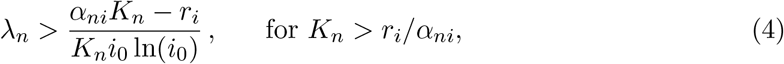

where *λ*_*n*_ represents an approximation for the minimal density-dependent microbiome transfer rate required, on average, for the establishment of the introduced population. Note that if the introduced population experiences a negative population growth even in the absence of competition with natives, Eq. 4 can be rewritten considering *α*_*ni*_ = 0 and *r*_*i*_ *<* 0 (see supplementary information, section D).

Eq. (4) tells us that increasing the number of introduced individuals *i*_0_ or the growth rate of the introduced population *r*_*i*_ increases the probability that the introduced species will establish, while increasing the competitive effect of natives on introduced individuals *α*_*ni*_ will decrease it (Fig. 2b). Increasing the carrying capacity *K*_*n*_ may increase or decrease the probability of establishment, depending on the strength of competitive interactions between natives and invaders and on the microbiome transfer rate. On the one hand, a large native population increases the probability that native microbes will be transfer to the introduced species in time to confer adaptation; on the other hand a large highly competitive native population may cause the extinction of the introduced population before microbial acquisition (cfr. Figs. S5b and S6).

### Scenario C: The presence of natives facilitates adaptation

Let us consider a certain patch in which a native population is present. Consider then that some highly competitive individuals of an invasive population are introduced to the patch. If introduced individuals have a higher competitive ability then natives, but are poorly adapted to local conditions, they may outcompete natives but remain present at low density after the invasion. If, however, invaders are conferred local adaptations through the acquisition of native microbes, their population may reach higher densities after the invasion. Thus, if the native population is displaced by the invaders before microbiome transfer can occur, invaders will remain in low numbers, otherwise their final population size will be larger (Fig. 3). Note that if the microbiome transfer rate is low, the increase in size of the invasive population is expected to be observed only after a lag time (Fig. S4b).

**Fig. 3:**
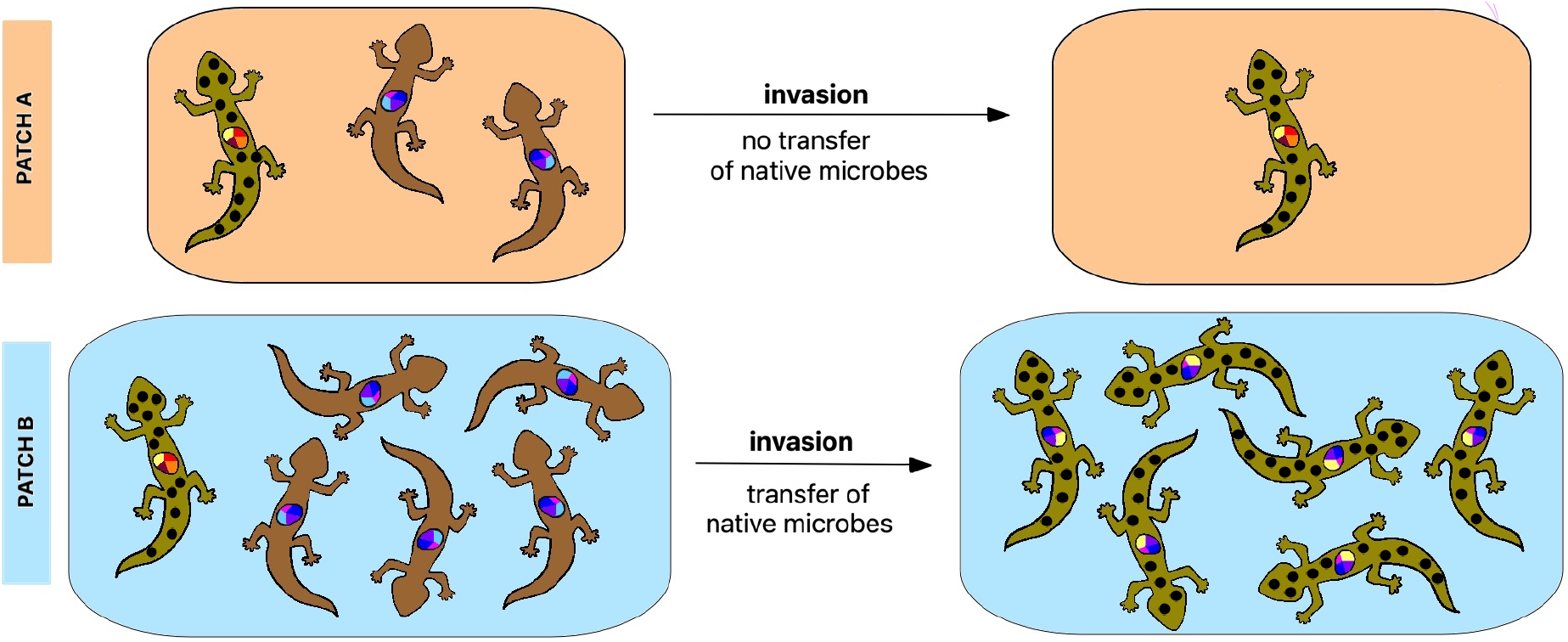
Conceptual representation of scenario C, of the mathematical results presented in Fig. S7. A native population (brown lizards) is competitively excluded by the introduction of a similar invasive species (green doted lizards). Patch A is smaller and presents a low density of natives, and the invaders displace the native population before the transfer of beneficial microbes from native to invaders can occur. In patch A, the invasive population remains therefore poorly adapted to the new environment, and in low density. Patch B presents a larger patch size and a higher density of natives, which increase the likelihood that native microbes will be transferred to the invaders before natives are driven to extinction. The acquisition of native microbes confers local adaptation to the invaders, and their population in patch B grows larger than in patch A. The findings represented in this figure are based on Eq. (5). The lizards here illustrate the possibility of such dynamics in species of animals or plants among which microbial sharing may occur. A conceptual representation of this figure is presented in Fig. 3.

The probability of acquiring microbes from natives will depend on the population densities of natives and invaders within a patch, on the nature of their interactions, and on the patch size. In the supplementary information (section E), we show that microbiome-mediated adaptation may occur when the minimal average density-dependent microbiome transfer rate *λ*_*n*_ satisfies

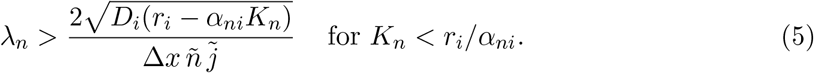

Eq. (5) tells us that the transfer of native microbes is more likely to occur when the average densities of natives 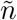 and invaders 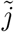 in a patch are large, when patch size Δ*x* is large, and when natives can slow down the growth of the invasive population through competition (i.e., *α*_*ni*_ is large enough, with *α*_*ni*_ *< r*_*i*_*/K*_*n*_). Under these circumstances, it will take longer for the invaders to outcompete natives, increasing their chance of acquiring native microbes and becoming locally adapted. Assuming that invaders disperse randomly within the patch, an increase in their intrinsic dispersal ability *D*_*i*_ will also lead to a faster displacement of natives, and thus to a lower probability of acquiring their microbiome.

## Discussion

### Scenario A The timing of microbiome acquisition affects invasion lag times

It is increasingly recognized that rapid evolution can alter invasion dynamics, where lags in biological invasion can emerge as a result of the time needed for evolutionary adaptation to take place in a new environment (Crooks, 2005; Whitney and Gabler, 2008). So far research has focused on genetic adaptation, while the evolutionary potential of non-genetic modes has begun to be explored only recently (Moran et al., 2021; Marin et al., 2020). The microbiome has been proposed as a non-genetic mode of conferring adaptability to host species (Kolodny et al., 2020; Henry et al., 2021), however the consequences of this adaptation for community dynamics have remained largely unexplored.

We propose that the adaptation of an introduced population to local conditions can be mediated by the acquisition of beneficial microbes which may have co-evolved locally with phylogenetically close natives. We also suggest that if the acquisition of microbes from the microbiome of native hosts increases the competitive ability of the introduced species, invasion can follow as a result. Thus, a lag in biological invasion may be observed because of the time required for an introduced species to acquire native microbes, where the duration of the lag time is determined by the rates of microbiome transfer between natives and invaders, and within the invasive population itself.

The idea that invaders may benefit from established mutualistic associations between native hosts and their microbes has already been formulated in plant ecology, where the establishment of an introduced plant and its expansion in a new range can be facilitated by the presence of pre-existing mycorrhizal networks (Dickie et al., 2017; Shipunov et al., 2008; Dawkins and Esiobu, 2016; Parepa et al., 2013). However, this concept is new to animal ecology, where inter-species horizontal transmission of mutualist microbes remains largely unexplored (Robinson et al., 2019; Bahrndorff et al., 2016). Research that links microbiome acquisition and host adaptation in animals is promising (Rennison et al., 2019; Fontaine et al., 2022; Kikuchi et al., 2012), but still in its infancy, and the problem of what is cause and what is consequence in host-microbiome relationships is unresolved in most cases. Going forward, it will be important to consider the intricate details of the mechanisms of host-symbiont interactions, both to better understand microbe’s role in fitness determination and in order to understand how specific these relations are.

There is no doubt that there are ample opportunities for microbial exchange to take place. Exchange of microbes can be brought about, for example, through predation of native individuals or through eating of their feces, a behaviour documented in invasive lizards (Norval et al., 2012a,b). Alternatively, environmentally mediated exchange could occur at sites of bathing in sand or water that are shared, e.g. between the invasive Indian myna (*Acridotheres tristis*) and many native species in the mynas’ sites of sunning, feeding grounds, and shelter. Our understanding of microbiome sharing among animal host species is currently limited (Bahrndorff et al., 2016), and the extent to which such exchange may result in the successful establishment of the natives’ microbes in the invasive species is unknown. Exploration of this topic is accordingly paramount to understanding the proposed scenario of microbiome-mediated adaptation.

The complementary tenet of this scenario is that inter-species sharing may provide significant adaptive value to the invasive species. As explained earlier, it seems highly likely that some microbial species that co-evolved over thousands of generations with a native host provide an adaptive value that may carry over to another host species that is related to it, such as a scenario in which a microbe that facilitates a certain carbohydrate’s breakdown in the gut of a native detritivore is picked up by a host species that belongs to the same ecological guild. These observations highlight multiple paths of empirical exploration that may provide major insights regarding microbiome-mediated adaptation in invasive species.

It is expected that if invaders can form novel mutualistic associations with microbes from the microbiome of natives, then hosts may also share pathogen strains (Dickie et al., 2017; Bahrndorff et al., 2016). In our work we choose to specifically focus on the case in which microbiome transfer has a positive impact on fitness, given that this scenario has received significantly less attention than the sharing of parasites or pathogens. The exchange of pathogens may also affect invasion dynamics, by reducing competitiveness in natives or in invaders. One prominent such example is in the case of the invasive grey squirrel, whose spread in Europe has been facilitated by infection of the native population of red squirrels with squirrelpox: a highly pathogenic disease carried by grey squirrels, which appear to be immune to it (Schuchert et al., 2014). We have recently outlined another interesting scenario along these lines as possibly having played a role in the spread of modern humans and the replacement of Neanderthals (Greenbaum et al., 2019). Future work may further consider how different combinations of mutualistic and parasitic/pathogenic interactions between microbes and a newly introduced host may affect species’ competitive dynamics and invasion success (Martignoni et al.).

### Scenario B The establishment of an introduced species is made possible by transfer of microbes from native species

In the previous section, we have already discussed how the presence of native mycorrhizal fungi in the soil may facilitate the establishment of newly introduced plants (Dickie et al., 2017; Becknell et al., 2021; Parepa et al., 2013). Experimental work is in progress to explore how specific microbiome can contribute not only to soil health and plant fitness, but also to animal reproductive success and in increasing their resilience against environmental stress (Peixoto et al., 2021; Comizzoli and Power, 2019), uncovering promising new venues for the successful management of reintroduced populations (Trevelline et al., 2019; Redford et al., 2012; Bahrndorff et al., 2016). Here we propose that in cases where the establishment of an introduced population is desired, such as the reintroduction of wildlife populations, the transfer of beneficial microbes from similar native species may increase establishment success by helping the introduced species to become better adapted to local conditions.

Founder populations have been often found to suffer from a lack of diversity that makes them more susceptible to demographic and environmental stochasticity (Drake and Lodge, 2006; Simberloff, 2009), and more likely to suffer from inbreeding depression (Drake and Lodge, 2006). The microbiome can influence the host phenotype in several ways (Fontaine et al., 2022; Kohl et al., 2014; Townsend et al., 2019), and phenotypic plasticity has been found to help founder populations with low genetic diversity to maintain high fitness (Richards et al., 2006; Davidson et al., 2011; Estoup et al., 2016). The extended phenotypic response provided by the acquisition of native microbes may therefore compensate for this lack in diversity and mediate the establishment success of small founder populations, particularly if native microbes are then efficiently transmitted among introduced individuals.

### Scenario C The presence of natives facilitates adaptation

Relatedness between invasive species and the recipient community have been found to be weak predictors of invasion success (Pantel et al., 2017; Leffler et al., 2014; Diez et al., 2008; Divíšek et al., 2018). On the one hand, similarities with natives may increase the likelihood that an invader’s traits will match the new environmental conditions. On the other hand, an invader may be more likely to suffer from direct competition with natives in such a case, due to niche overlap. Here we propose that, non-intuitively, invasion may be *facilitated* by the presence of co-occurring native species if the acquisition of beneficial pre-adapted microbes from the microbiome of natives can boost invaders’ fitness.

Particularly, even when invaders are superior competitors, the acquisition of native microbes may confer local adaptations to an invasive population and facilitate its population growth and spread. On the other hand, if invaders displace natives before being able to acquire their microbiome, invaders may fail to adapt to the new environment and remain localized in certain patches or regions. Eventually, due to being poorly adapted, environmental disturbance may cause their disappearance after what seemed to be a successful invasion, a phenomenon that has been observed in several cases, some of them not fully understood (Simberloff, 2013). Such an example is the spread of Indian palm squirrels (*Funambulus pennati*) in Israel and their eventual disappearance, perhaps because of a cold spell during winter (Yom-Tov, 2013).

Our hypothesis could be tested experimentally in controlled conditions that emulate invasion scenarios, comparing the invaders’ fitness when faced with local conditions, with and without exposure to native species that may act as potential microbiome sources for local adaptation. Perhaps more interestingly, it may also be explored in invasive species that were introduced independently multiple times to sites which are disconnected. An example of such introduction is that of the marbled crayfish (*Procambrus virginalis*) in different places around the world. This species has been introduced in some cases to multiple wetlands in the same region that are characterized by similar environmental conditions, but that differ in the native crustacean hosts that occur in them and that may function as native microbiome ‘donors’. Comparing the crayfishes’ fitness and invasive success among these sites, and linking them to the microbiome site composition of the native and invasive crustaceans, may thus be highly informative. We have recently detected and have begun to study such a situation in Israel, where the marbled crayfish was recently detected at several sites (a report of this set of invasions and their eradication attempts is in preparation).

The number of studies comparing the microbiome of native and invasive species in plants is growing rapidly (Coats and Rumpho, 2014; Aires et al., 2021), but only a few studies have focused on comparing the microbiome of native and invasive animals (Chiarello et al., 2022; Santos et al., 2021). In a recent study, Chiarello et al. (2022) found that native mussels shared a substantial fraction of their microbiome with the co-occurring invasive species *Corbicula fluminea*, indicating that invasive mussels may host microbial communities that are obtained locally. Additionally, a few more studies have compared the microbiome of invaders in their native and invasive range (Cardoso et al., 2012; Bansal et al., 2014), or in the population core and at the edges of their expansion range (Dragičević et al., 2021; Wagener et al., 2021). We suggest that such studies are necessary for the understanding of the possible importance of microbiome-mediated adaptation in general, as well as for testing the proposed hypothesis of adaptive microbiome pickup from native hosts as a mode of rapid adaptation. In a rapidly changing world in which connectivity and opportunities for the spread of invasive species are consistently increasing, these may turn out to be key to understanding and predicting species’ invasion success, and in turn to considering the mode and timing of mitigation efforts.

## Conclusion

The need for developing theoretical frameworks to predict invasive potential when invaders evolve in their environment has been highlighted in several instances (Coutts et al., 2018; Whitney and Gabler, 2008), however this call has largely remained unanswered. Here we present a mathematical model that sheds light on possible dynamics occurring if invaders evolve after their introduction, and we focus on the situation in which evolution is driven by the transfer of beneficial microbes from the microbiome of similar co-occurring native species. Our work presents a simple framework which sets the basis for broadening the conceptualization of microbiome-mediated dynamics, and opens the door to further theoretical exploration and scientific discoveries.

## Acknowledgement

MM is founded by the Azrieli Foundation. OK and MM are founded by the Israel Science Foundation (ISF) (grant number 1826/20), the Gordon and Betty Moore Foundation, and the United States-Israel Binational Science Foundation (BSF).

## Supplementary information

### A Two-species competition model

Consider the competitive Lotka-Volterra equations to describe the competitive dynamics between a native population (*N*) and an introduced population (*I*_0_):

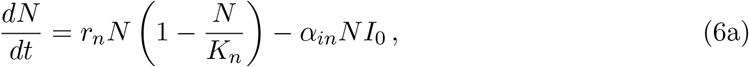

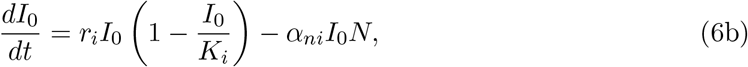

where *r*_*n*_ and *r*_*i*_ are the intrinsic growth rates of the native and introduced populations, *K*_*n*_ and *K*_*i*_ are their carrying capacities, and *α*_*in*_ and *α*_*ni*_ quantify the competitive effect of the introduced species on natives, and viceversa. A description of model parameters is provided in Table 1.

**Table 1:**
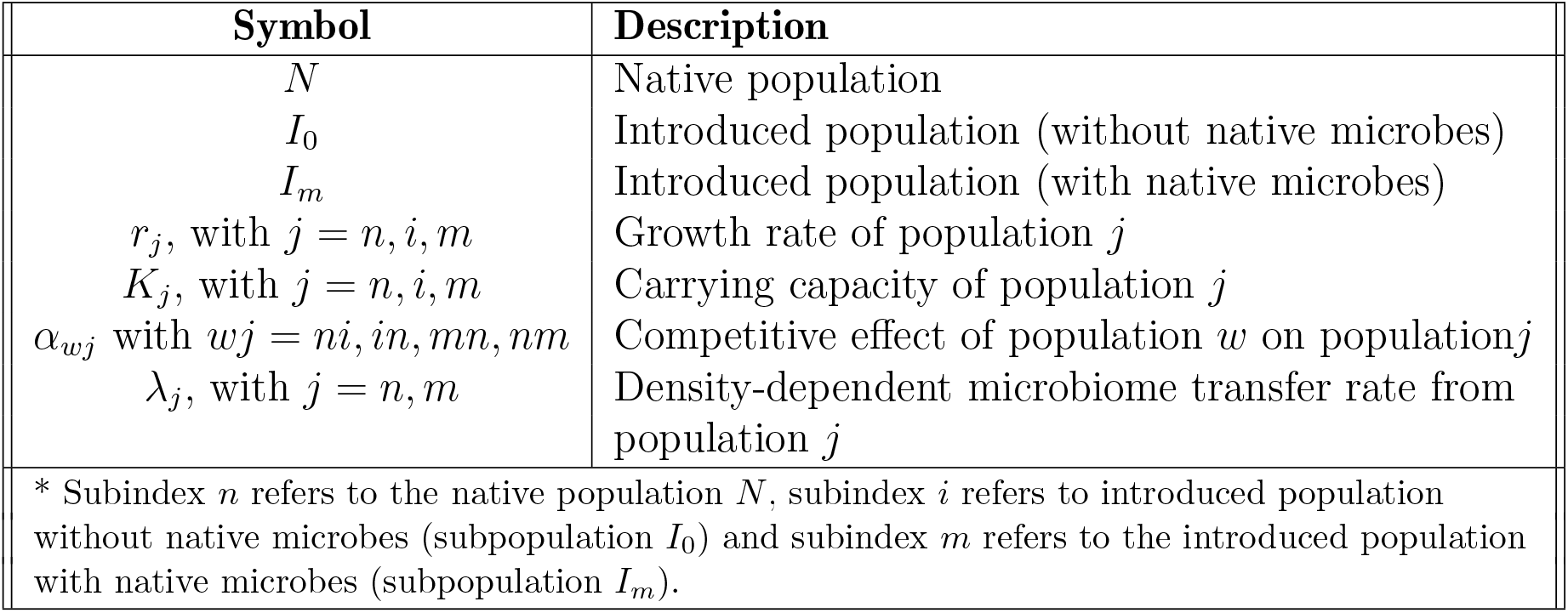
Brief description of the variables and parameters of the system of equations in (1).

| Symbol | Description |
| --- | --- |
| $N$ | Native population |
| $I_0$ | Introduced population (without native microbes) |
| $I_m$ | Introduced population (with native microbes) |
| $r_j$, with $j = n, i, m$ | Growth rate of population $j$ |
| $K_j$, with $j = n, i, m$ | Carrying capacity of population $j$ |
| $\alpha_{wj}$ with $wj = ni, in, mn, nm$ | Competitive effect of population $w$ on population $j$ |
| $\lambda_j$, with $j = n, m$ | Density-dependent microbiome transfer rate from population $j$ |
| * Subindex $n$ refers to the native population $N$, subindex $i$ refers to introduced population without native microbes (subpopulation $I_0$) and subindex $m$ refers to the introduced population with native microbes (subpopulation $I_m$). | |

Linear stability analysis and phase plane analysis (Kot, 2001) show that the dynamics of the system of equations in (6) can result in the four different scenarios described below. The phase planes corresponding to each of these scenarios are shown in Fig. S1.

#### (a) Coexistence of natives and introduced species

Coexistence of the native and introduced populations is observed when growth rates and competition rates between natives and introduced species are low, and their carrying capacities are high (i.e., when *K*_*n*_ *< r*_*i*_*/α*_*ni*_ and *K*_*i*_ *> r*_*n*_*/α*_*in*_, see Fig. S1a). Under this scenario, the coexistence steady state (*N* ^*\**^, 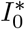) is stable, with

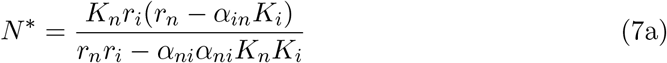

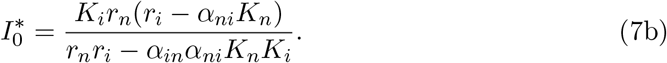

Thus, if competition between native and introduced species if low, species coexists at an equilibrium value that is lower than their respective carrying capacity.

#### (b) Competitive exclusion (bistability)

When both natives and introduced species are strong competitors, such that *K*_*i*_ *> r*_*n*_*/α*_*in*_ and *K*_*n*_ *> r*_*i*_*/α*_*ni*_, coexistence cannot occur and competitive exclusion of natives or introduced species is observed. In this case, both steady states (*N* ^*\**^, 0) and (0, 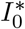), with *N* ^*\**^ = *K*_*n*_ and 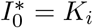, are stable, and which species will competitively exclude the other will depend on model parameters (determining the size of the basin of attraction of each of the steady states), and on the initial conditions (Fig. S1b).

#### (c) Competitive exclusion of the introduced species

If natives are superior competitors, only the steady state (*N* ^*\**^, 0) with *N* ^*\**^ = *K*_*n*_ is stable, and natives will competitively exclude the introduced species (Fig. S1c). This scenario may occur if the carrying capacity and growth rate of the introduced species, and if the competitive effect of natives on the introduced population are low, while the carrying capacity and growth rate of natives and the competitive effect of introduced species on natives are high, such that *K*_*n*_ *> r*_*i*_*/α*_*ni*_ and *K*_*i*_ *< r*_*n*_*/α*_*in*_.

#### (d) Competitive exclusion of natives

If the introduced species is a superior competitor, only the steady state (0, 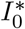) with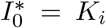 is stable and the introduced population will competitively exclude natives (Fig. S1d). This scenario may occur if the carrying capacity and growth rate of the introduced species, and its competitive effect on natives are high, while the carrying capacity and growth rate of natives, and their competitive effect on invaders are low, such that *K*_*i*_ *> r*_*n*_*/α*_*in*_ and *K*_*n*_ *> r*_*i*_*/α*_*ni*_.

**Fig. S1:**
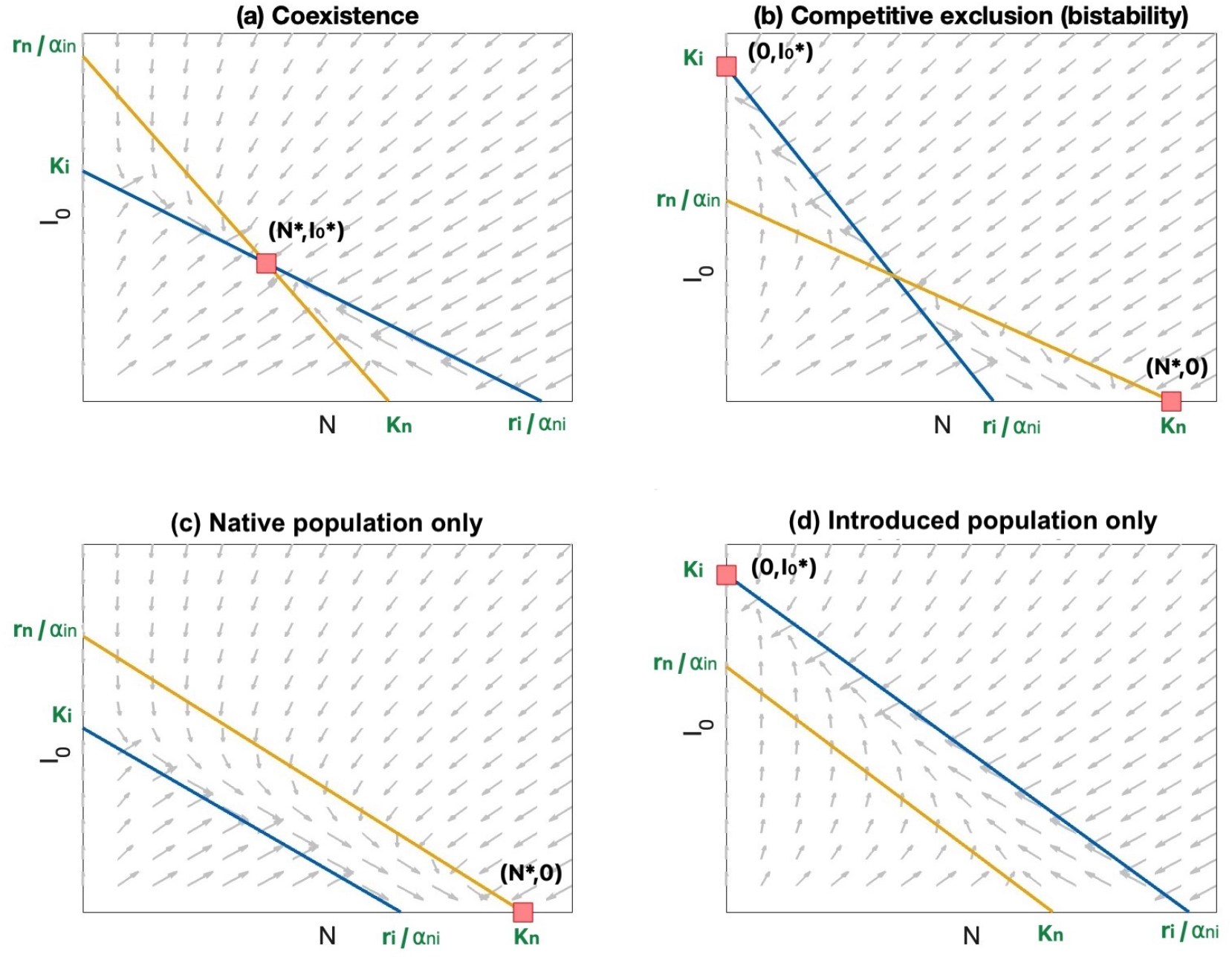
Phase planes of the system of equations in (6). Nullclines are represented in orange (*dN/dt* = 0) and blue (*dI*_0_*/dt* = 0). The horizontal axis represents the native population, while the vertical axis represents the introduced population. Stable steady states are represented with a red square. We observe that depending on the competitive effect that natives and introduced species have on each other, their carrying capacity, and their growth rates, different scenarios can be observed, namely (a) coexistence between the native and the introduced populations, (b) competitive exclusion of one of the two populations, where which population survives will depend on model parameters and initial conditions, (c) competitive exclusion of the introduced population, and (d) competitive exclusion of natives.

### B Dynamics of competition and microbiome transfer

The transfer of beneficial microbes from the native to the introduced population can lead to an increased competitive ability of the introduced population. We model this scenario by splitting the introduced population *I* in two subpopulation, i.e., the subpopulation without native microbes *I*_0_ and the subpopulation with native microbes *I*_*m*_ (Eq. (1)). Once the first microbiome transfer event from natives to introduced individuals occurred, a new subpopulation *I*_*m*_ is created. We model the increase in competitive ability due to the presence of native microbes by allowing the *I*_*m*_ subpopulation to have for example a higher carrying capacity than *I*_0_ (i.e., *K*_*m*_ *> K*_*i*_) or a higher growth rate (i.e., *r*_*m*_ *> r*_*i*_). The biological reasons for this choice are explained in the main manuscript.

Looking at the phase plane of the two-species competition model (Fig. S1) can help us visualize the impact of creating a new subpopulation with superior competitive ability on the competitive dynamics. We would like to use a phase plane representation to understand the impact that microbiome transfer can have on the competitive dynamics in scenarios A, B and C, described in the main manuscript.

#### Scenario A The timing of microbiome acquisition affects invasion lag times

Prior to the transfer of native microbes, the introduced species coexists in low density with a much more abundant native population. Microbiome transfer from native to introduced individuals causes an increase in the carrying capacity of the introduced population. This increase constitutes a competitive advantage for the introduced population, that eventually leads to the competitive exclusion of natives, or to a significantly reduction in their population size (Fig. S2, scenario A).

#### Scenario B The establishment of an introduced species is made possible by transfer of microbes from native species

Natives are competitively superior to the introduced species, and if microbiome transfer from natives does not occur the introduced population would fail to establish. If microbiome transfer occurs before the extinction of the introduced population, it may lead to an increase in the carrying capacity, and eventually in the growth rate, of the introduced species, and the consequent competitive exclusion of natives (Fig. S2, scenario B).

#### Scenario C The presence of natives facilitates adaptation

The introduced species is competitively superior than natives, however despite its high competitive ability, it has a low carrying capacity and growth rate. If microbiome transfer does not occur, the introduced population displaces the native population, but its population size remains small. Microbiome transfer increases the carrying capacity of the introduced population, and allows it to reach a higher population size after the displacement of natives (Fig. S2, scenario C).

**Table S1:** Default parameters used to simulate scenarios A, B and C. The corresponding phase planes are provided in Fig. S2.

| Parameter | Scenario A | Scenario B | Scenario C |
| --- | --- | --- | --- |
| $K_n$ | 100 | 90 | 100 |
| $K_i$ | 20 | 60 | 20 |
| $K_m$ | (b) 70 (c) 90 | (b) 60 (c) 90 | 80 |
| $r_n$ | 1.5 | 1.5 | 1.5 |
| $r_i$ | 1.5 | 1.5 | 1.5 |
| $r_m$ | 1.5 | 2.5 | 1.5 |
| $\alpha_{ni}$ | 0.01 | 0.02 | 0.01 |
| $\alpha_{in}$ | 0.02 | 0.02 | 0.2 |
| $\alpha_{nm}$ | 0.01 | 0.02 | 0.01 |
| $\alpha_{mn}$ | 0.02 | 0.02 | 0.2 |

**Fig. S2:**
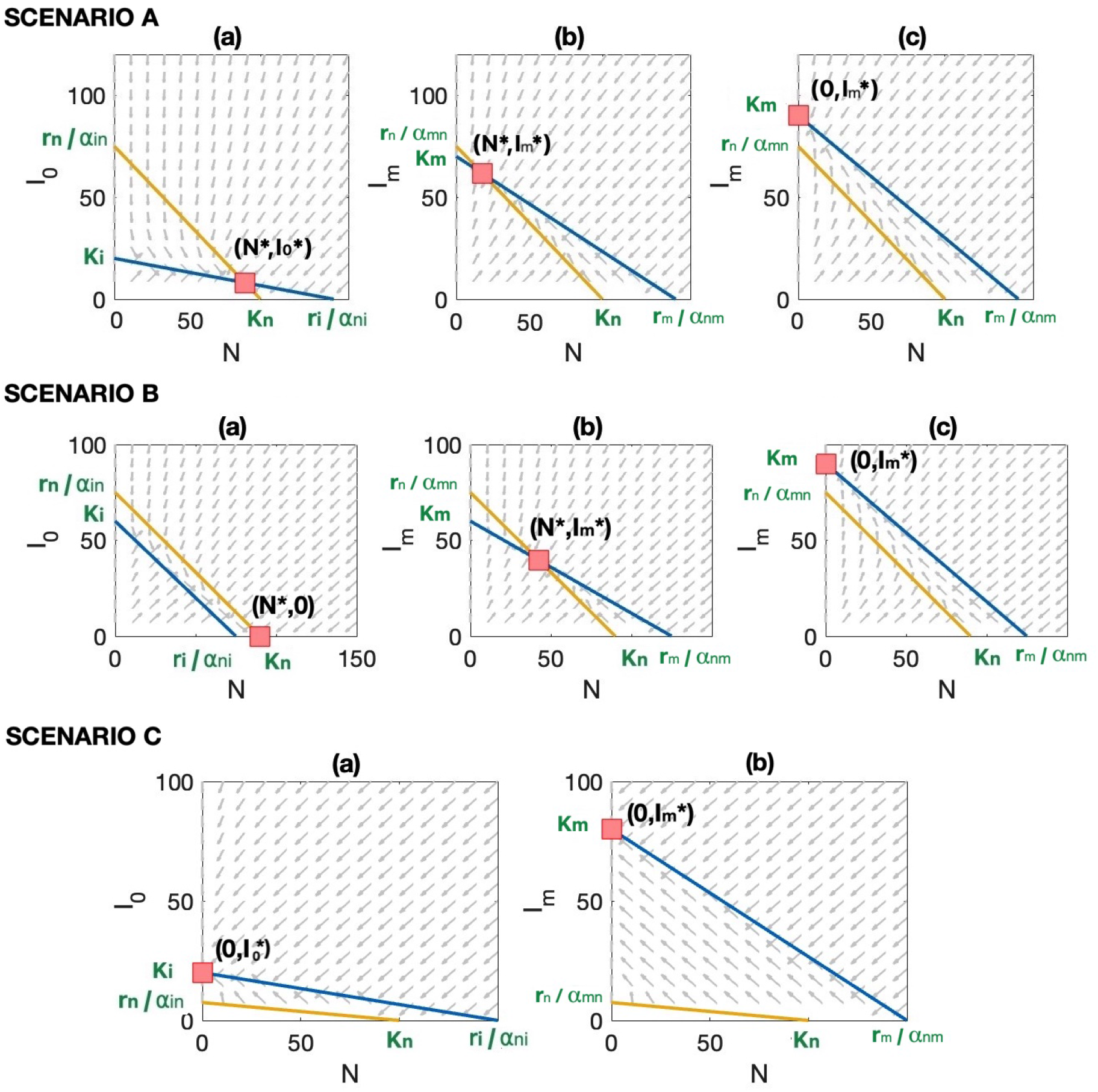
Phase plane representation of the two-species competition model of Eq. (6), where a native population competes with an introduced population that has not acquired native microbes (*I*_0_), or that has acquired native microbes (*I*_*m*_). Scenario A: (a) Coexistence between the native and introduced populations is observed prior to the transfer of native microbes, where the introduced population is present in small numbers. Microbiome transfer leads to an increase in the carrying capacity of the introduced population (from *K*_*i*_ to *K*_*m*_ *> K*_*i*_), and a consequent (b) reduction or (c) extinction of the native population. Scenario B: (a) the native population is competitively superior to the introduced population. Microbiome transfer causes (b) an increase in the growth rate (from *r*_*i*_ to *r*_*m*_ *> r*_*i*_), and eventually (c) an increase in the carrying capacity of the introduced population (from *K*_*i*_ to *K*_*m*_ *> K*_*i*_), which leads to the (b) reduction or (c) extinction of the native population. Scenario C: (a)The introduced population is competitively superior, and its introduction leads to the exclusion of the native population. (b) Microbiome transfer leads to an increase in the carrying capacity of the introduced population (from *K*_*i*_ to *K*_*m*_ *> K*_*i*_). The parameters used to simulate these three scenarios are provided in Table S1.

## C Stochastic realizations and deterministic approximation

The ordinary differential equation system of Eq. (1) includes two Poisson random variables, namely Λ_*n*_ and Λ_*m*_, defined in Eqs. (2) and (3). These two random variables representing microbiome transfer from natives to introduced individuals (Λ_*n*_) and among introduced individuals (Λ_*m*_), where the expected population increase of *I*_*m*_ due to microbiome transfer 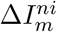 is

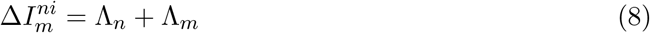

Conditioning on 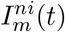, we have that:

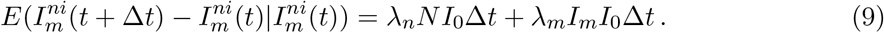

Dividing Eqs. (9) by Δ*t* and letting Δ*t →* 0, we obtain:

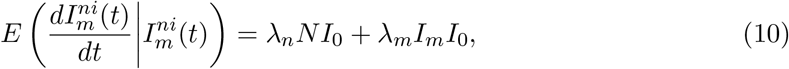

what corresponds to the the deterministic version of the model of Eq. (1). Thus as long as *E*(Λ_*n*_) and *E*(Λ_*m*_) are large enough, we expect the mean average of a large number of stochastic realizations of Eq. (1) to approach the deterministic solution.

In Fig. S3 we plot 500 stochastic realizations of Eq. (1), and compare the mean average of these realizations with the corresponding deterministic solution of Eq. (1), for which the random variables Λ_*n*_ and Λ_*m*_ are substituted by their expected values. We can see that for scenario A, for *λ*_*n*_ and *λ*_*m*_ large enough, the mean average of a large number of realizations approaches the deterministic solution.

**Fig. S3:**
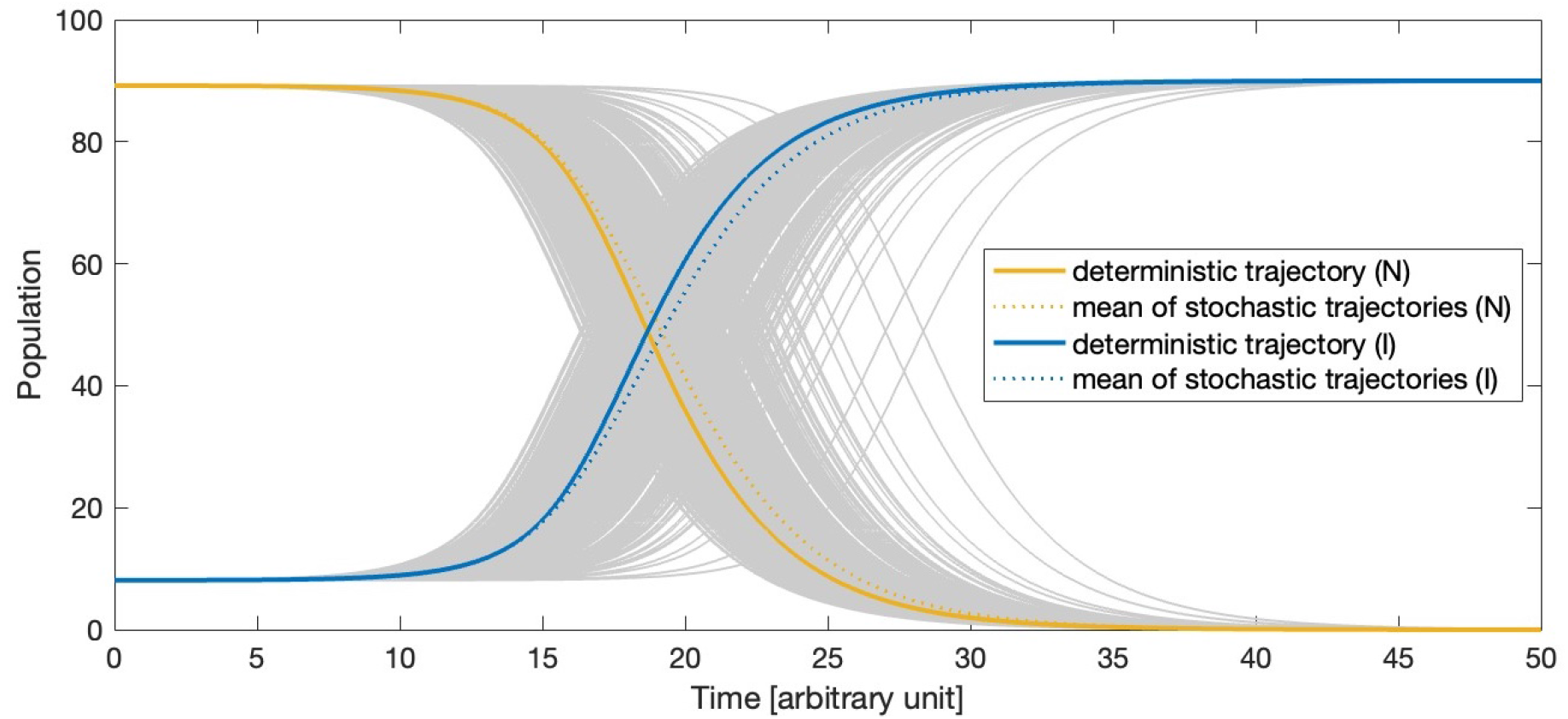
Stochastic realizations of Eq. (1) (grey curves) are computed under the scenario A, in which the introduced species coexists with the native population prior to the transfer of native microbes (cfr. Fig. 1a). The mean of 500 realizations is represented as dotted line and the deterministic solution is represented as a solid line, for the native (orange) as well as for the introduced population (blue). Note that the mean of a large number of stochastic realizations approaches the deterministic solution.

### D Establishment in the presence of natives

Consider the situation in which the native population is competitively superior to the introduced population *I*_0_ (Fig. S2, scenario B). In this case, microbiome transfer can contribute to increase the competitive ability of the introduced population, and rescue it from extinction (Fig. S4a).

We are interested in deriving an approximation for the minimal microbiome transfer rate needed to avoid the extinction of the introduced population. For this purpose, we look at the equation describing the growth rate of the introduced population *I*_0_ prior to the first microbiome transfer event, namely Eq. (1b), for *I*_*m*_ = 0:

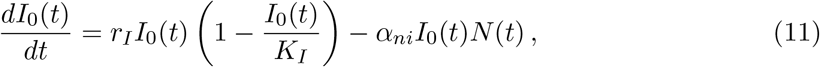

Let us assume that only a small number of individuals *i*_0_ is introduced, such that *I*_0_ *≪ K*_*I*_ and *N* (*t*) = *K*_*n*_. We obtain:

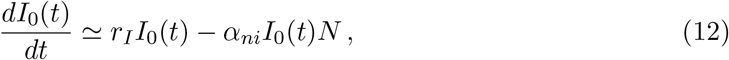

which has solution

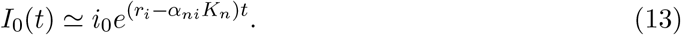

We know that for the scenario considered *K*_*n*_ *> r*_*i*_*/α*_*ni*_ (Fig. S2, scenario B(a)), i.e., *r*_*i*_ − *α*_*ni*_*K*_*n*_ *<* 0 and the population *I*_0_(*t*) declines exponentially. We define *t*_0_ as the time needed to drive the introduced population to extinction. Note that *I*_0_(*t*), as approximated in Eq. (13), never reaches zero, while in the two-species competition model (Eq. (6)) the native population *N* competitively exclude *I*_0_. This is a limitation due to considering *N* (*t*) as a constant. We approximate therefore *t*_0_ as the time needed for the introduced population to reduce to a single individual:

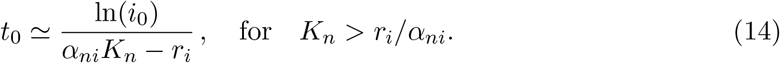

Eq. (14) can serve as a good approximation to understand how model parameters can affect the minimal microbiome transfer rate needed to rescue the introduced population from extinction. Note that if we consider that the introduced population has a negative population growth even in the absence of competition (i.e., *r*_*i*_ *<* 0, and Eq. (12) becomes *dI*_0_*/dt* = *r*_*i*_*I*_0_), the introduced population will decline even in the absence of a native population.

The expected value of the microbiome transfer rate between native and introduced individuals (Eq. (2)) is

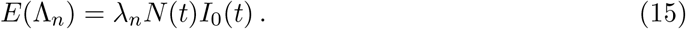

We define *t*_*n*_ as the average time needed for the first microbiome transfer event to occur. Assuming a constant native population *K*_*n*_ and an initial introduced population *i*_0_, we obtain a lower and a upper bound for *t*_*n*_ during the exponential decay of population *I*_0_(*t*) from *i*_0_ to 1, namely:

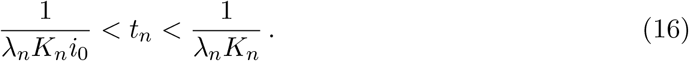

Thus, the larger the number of introduced individuals, the shorter the expected time *t*_*n*_ till the occurrence of the first microbiome transfer event. A reduction in the size of the introduce population *I*_0_(*t*) also implies a reduction in the likelihood of transferring native microbes, i.e., an increase in *t*_*n*_.

Using Eqs. (14) and (16) we can understand how the minimal density-dependent microbiome transfer rate *λ*_*n*_ needed to avoid extinction of the introduced population relates to other model parameters, i.e., the situation in which *t*_*n*_ *< t*_0_. We obtain the following approximation for the lower and upper bounds for the minimal density-dependent microbiome transfer rate *λ*_*n*_:

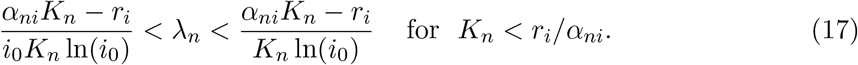

Eq. (17) holds best when *i*_0_ *≪ K*_*i*_ and when the native population is little affected by the presence of an introduced species. Eq. (17) be used to identify key parameters affecting the competition dynamics under scenario B, and determine how their variation affects the establishment success of the introduced population.

Plots of the upper bound of the minimal *λ*_*n*_ as a function of the size of the introduced population *i*_0_, the growth rate of the introduced population *r*_*i*_, the competitive effect of the native on the introduced population *α*_*ni*_ and the carrying capacity of the native population *K*_*n*_ are shown in Fig. S5. Numerical simulations investigating the probability of establishment as a function of *λ*_*n*_ and *i*_0_ are shown in the main manuscript (Fig. 2b). Note that increasing the initial population of the introduced species (parameter *i*_0_), or increasing its growth rate *r*_*i*_ leads to population establishment for a lower microbiome transfer rate *λ*_*n*_ (Fig. S5 (a) and (d)). A higher microbiome transfer rate *λ*_*n*_ is required for the establishment of the introduced population when the size of the native population is large (higher carrying capacity *K*_*n*_) or when the competitive effect of the native on the introduced population *α*_*ni*_ is high (Fig. S5 (b) and (c)).

Note that if the introduced population experiences a population decline even in the absence of competition with natives, the denominator *α*_*ni*_*K*_*n*_ − *r*_*i*_ of Eq. (17) can be substituted by −*r*_*i*_, with *r*_*i*_ *<* 0 representing the rate of decline in the size of the introduced population, and Eq. (17) becomes

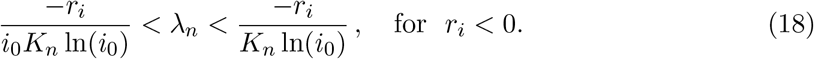

In this case, the upper bound of the minimal *λ*_*n*_ increases as a function of the carrying capacity of the native population *K*_*n*_ (Fig. S6), while the influence of other parameters will remain similar. Indeed, the larger the native population, the higher the probability that introduced individuals will acquire native microbes, and thus in the absence of competition a large native population will only be beneficial for the introduced population.

**Fig. S4:**
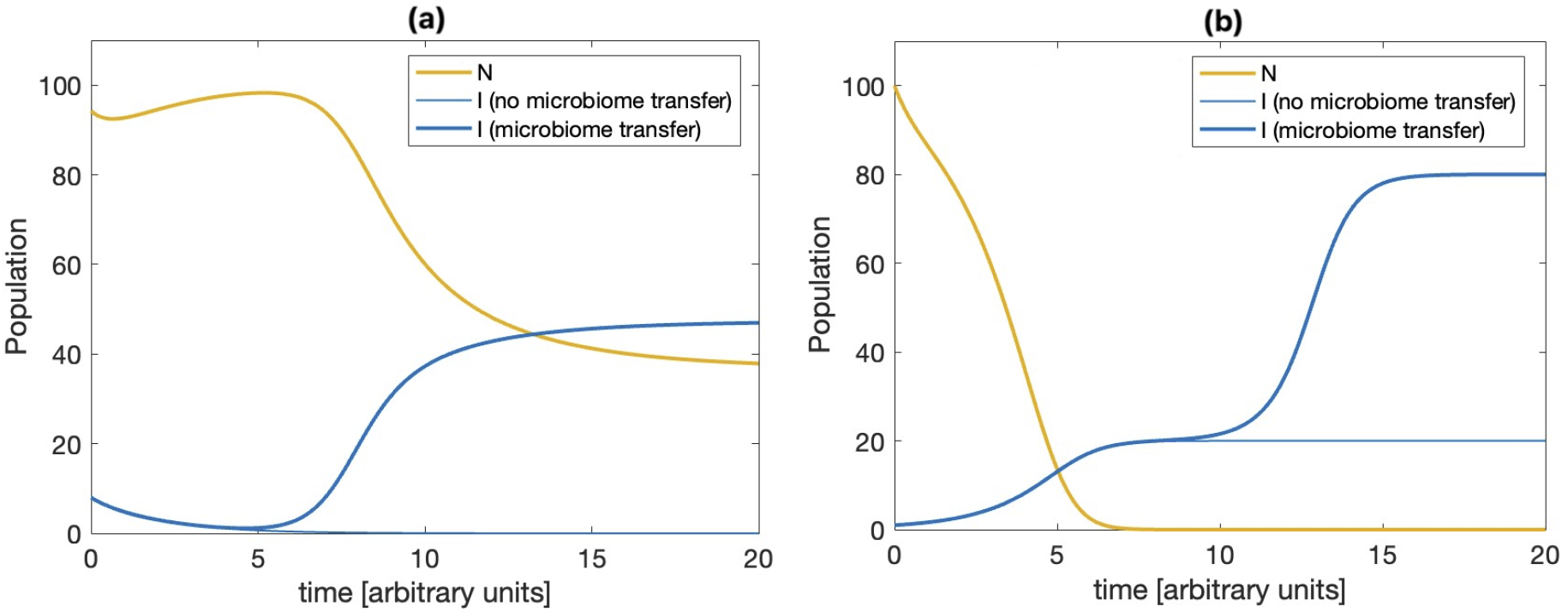
Growth of the native (*N*, orange curve) and introduced population (*I*, blue curve) over time when microbiome transfer from natives does not occur (thin blue curve), and when it does (thick blue curve). (a) When natives are superior competitors (scenario B) microbiome transfer can facilitate the establishment of an introduced species. If the number of introduced individuals is large enough, and if the rate of microbiome transfer is small, the establishment of the introduced population may occur after a lag time. (b) If the invaders are superior competitors, natives are displaced by their introduction (scenario C). The transfer of beneficial microbes can facilitate the rapid adaptation of invaders and increase their carrying capacity. If the rate of microbiome transfer is low, the increase in carrying capacity may be observed after a lag time.

**Fig. S5:**
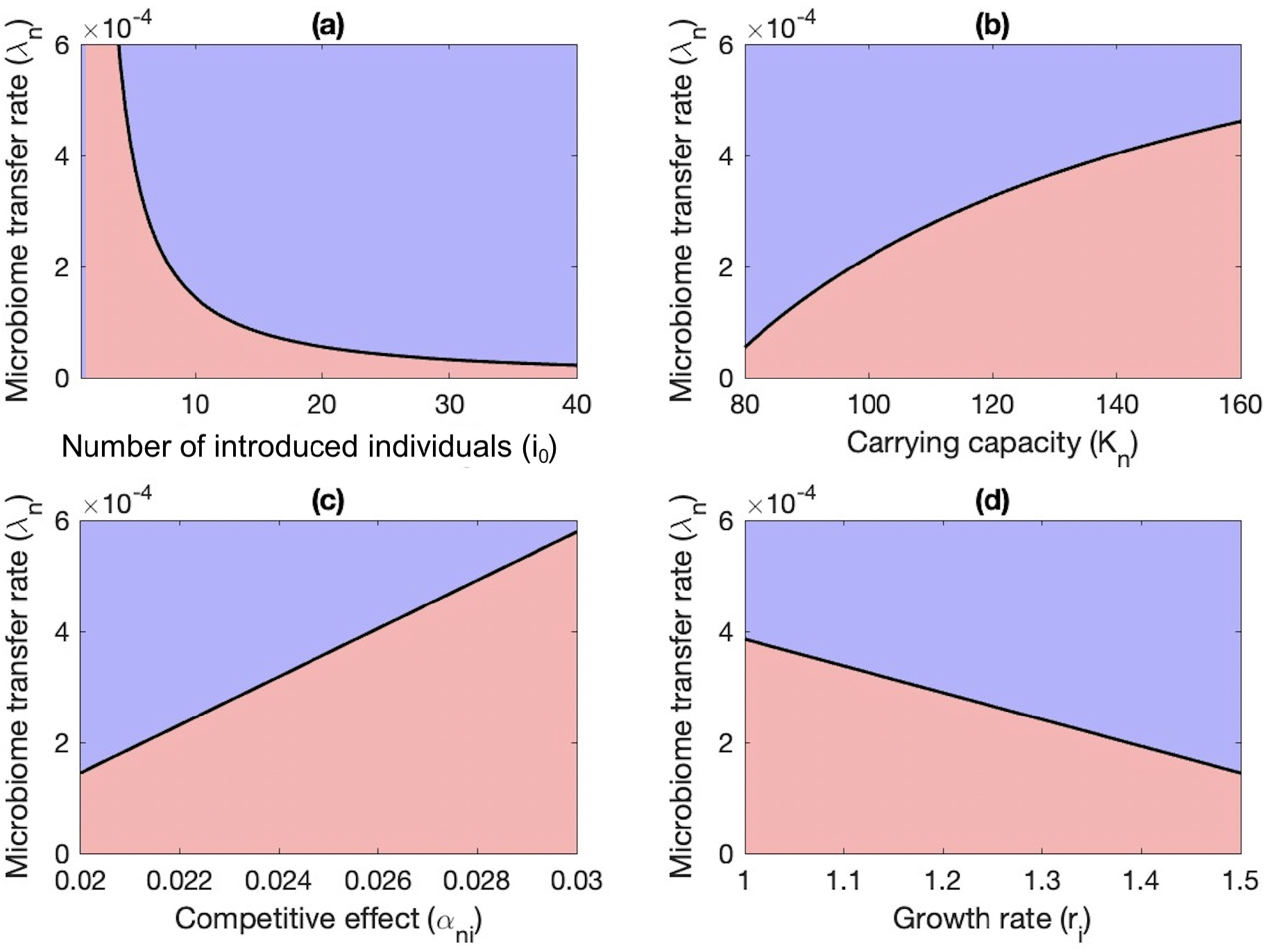
Upper bound of the minimal density-dependent microbiome transfer rate (*λ*_*n*_) needed for the establishment of an introduced population, when the native population is competitively superior (Eq. (17)). Parameter *λ*_*n*_ is plot as a function of (a) the initial size of the introduced population *i*_0_, (b) the size of the native population, assumed to be at carrying capacity *K*_*n*_, (c) the competitive effect of natives on introduced species *α*_*ni*_, and (d) the growth rate of the introduced population *r*_*i*_. In all figures, the blue region represents establishment of the introduced species, while red region represents its extinction.

**Fig. S6:**
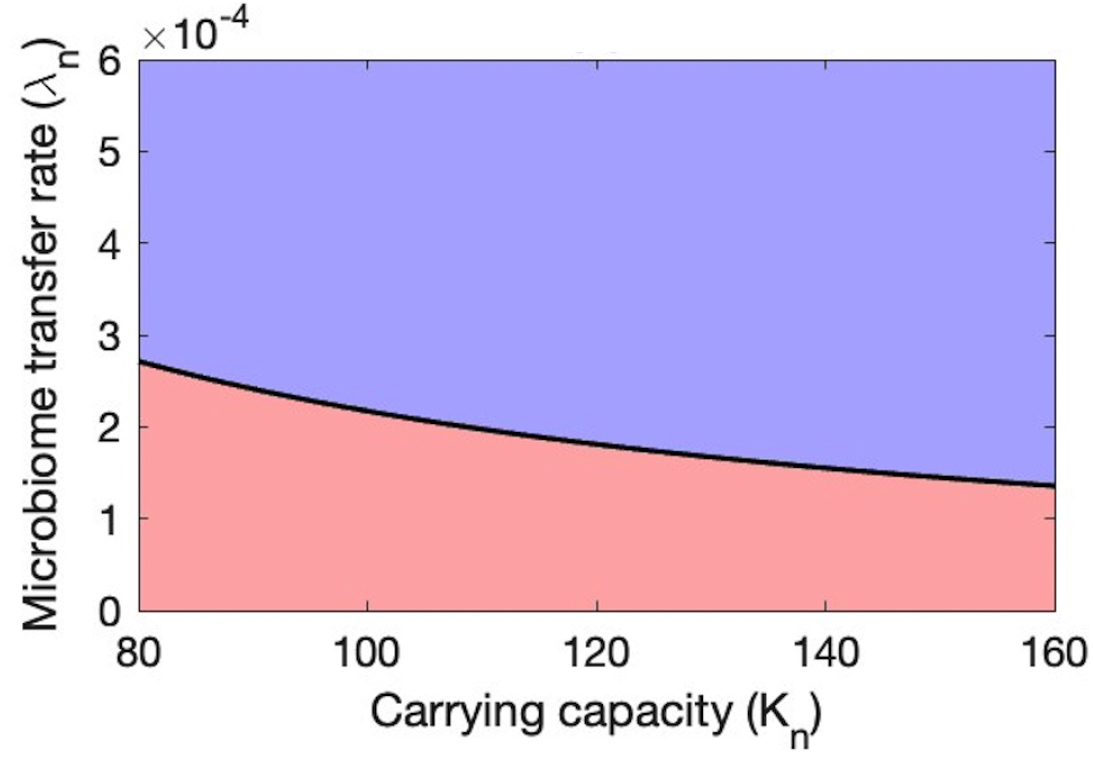
Upper bound of the minimal density-dependent microbiome transfer rate (*λ*_*n*_) needed for the establishment of an introduced population, when the introduced population experienced a population decline even in the absence of competition with natives (Eq. (18)). Parameter *λ*_*n*_ is plot as a function of the carrying capacity *K*_*n*_. The blue region represents establishment of the introduced species, while red region represents its extinction.

### E Speed of invasion

We consider the situation in which introduced individuals are competitively superior, and their introduction leads to the exclusion of natives (Fig. S2, scenario C). In our simulations, microbiome transfer from native to introduced individuals facilitates the adaptation of the introduced population, and we are interested in understanding under which circumstances microbiome transfer can occur before the displacement of natives. Would microbiome transfer not occur in time, the introduced population still displaces native species, but remains poorly adapted in the environment (Fig. 3 and Fig. S4).

For this purpose, we consider a spatially explicit version of the two species competition model presented in A. In this case, the two species represent a population of natives *N*, and a competitively superior introduced population *I*_0_ that can disperse in a given homogeneous one-dimensional landscape *x*. We consider that the introduced population disperses randomly, and we quantity its dispersal ability by the diffusion coefficients *D*_*i*_. Diffusion of *I*_0_ causes the subsequent displacement of natives. We write:

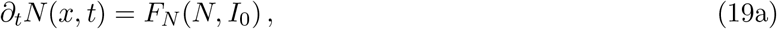

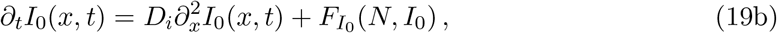

with

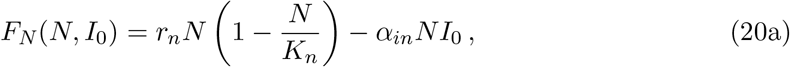

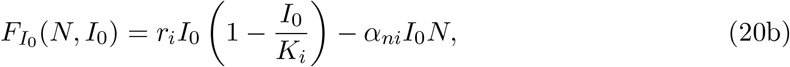

as defined in Eq. (6), where model parameters are given in Table 1. The population densities of natives and introduced individuals at each time *t* and location *x*, i.e., variables *N* (*x, t*) and *I*_0_(*x, t*), are given by the solutions to Eq. (19).

#### E.1 Traveling wave analysis

We would like to derive an approximation for the speed at which a competitively superior introduced population *I*_0_ will displace a native population *N* (Fig. S2, scenario C). To tackle this problem, we look at travelling wave solutions of Eq. (19), which are particular solutions describing the invasion at a constant speed *c* of the steady state (*N* ^*\**^, 0), for which only natives are present, by the steady state (0, 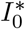), for which only the introduced species is present. We assume therefore solutions to Eq. (19) to be of the form *N* (*x, t*) = *n*(*x* − *ct*) = *n*(*z*) and *I*_0_(*x, t*) = *j*(*x* − *ct*) = *j*(*z*) for an unknown speed *c* ∈ ℝ. By replacing these expressions into (19) we obtain:

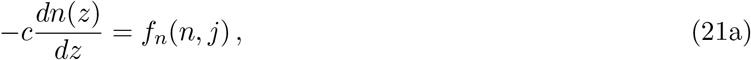

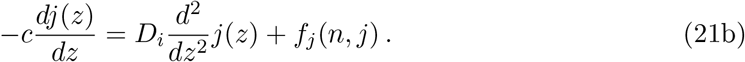

We consider that initially a native population is at its carrying capacity *K*_*n*_, and that successively the native population is displaced by the introduced population, which is advancing at constant speed *c* from left to right in the *z*-domain. This situation can be modelled by assuming that at the right of the domain (i.e., ahead of the wave, for *z→* +*∞*) a native population is present at its steady state (*n*^*\**^, 0), with *n*^*\**^ = *K*_*n*_, while on the left side of the domain (i.e., behind the wave, for *z →* −*∞*), only the introduced population *j* is present at steady state (0, *j* ^*\**^), with *j* ^*\**^ = *K*_*i*_. Thus the boundary conditions can be expressed as:

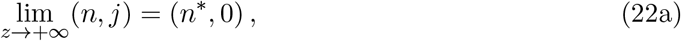

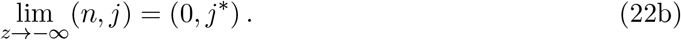

Under scenario C, we know that the steady state (*n*^*\**^, 0) is unstable, while (0, *j*^*\**^) is stable. When a stable and an unstable steady states are present, the stable steady state will invade the unstable one at constant speed *c*. In this case, there is a monostable traveling wave solution (mimicking biological invasion) and one may expect an estimate of the minimal speed of propagation *c* using the linearized problem around the unstable steady state.

To find the minimal speed of propagation *c* we define the variable *u* = *j*^*′*^, such that the system of equations in (21) can be rewritten as a system of first order ordinary differential equations:

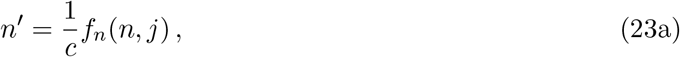

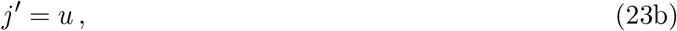

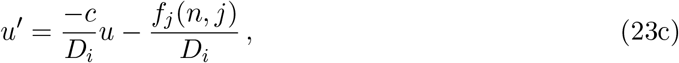

with Jacobian

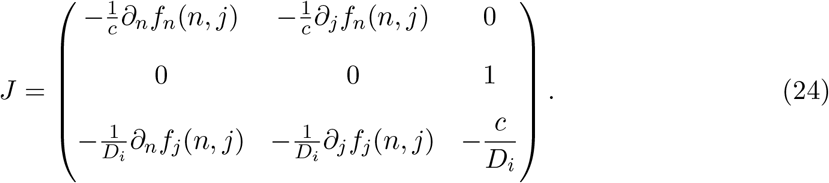

When computing the Jacobian *J* around the unstable steady state (*n*^*\**^, 0) with *n*^*\**^ = *K*_*n*_, and using Eqs. (20a) and (20b), we obtain:

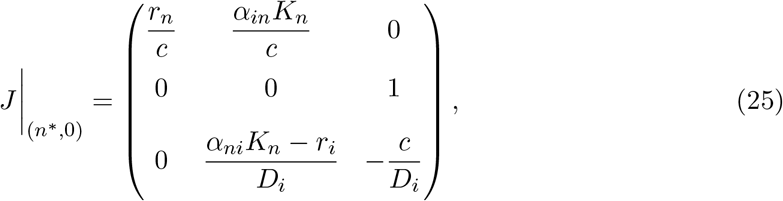

with eigenvalues *λ* corresponding to the solutions of

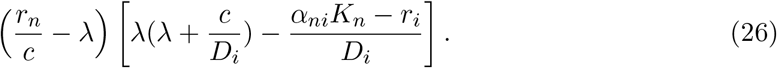

Thus,

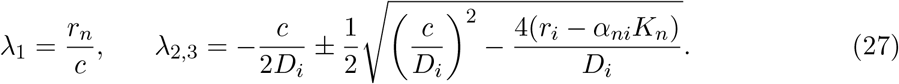

We know that, in scenario C, *r*_*i*_ −*α*_*ni*_*K*_*n*_ *>* 0. Additionally, we are only interested in solutions that are bounded below by zero, as *n*(*x, t*), *j*(*x, t*) *>* 0. For this, we require (*n*^*\**^, 0) to be a stable focus. In other words, we require

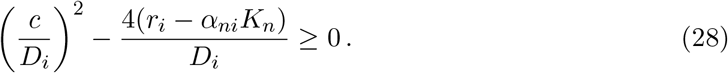

Hence, one can conclude that a lower bound for the propagation speed *c*^*\**^ *≤ c* is given by

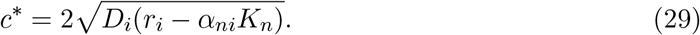

The speed of invasion *c* increases therefore with increasing dispersal ability of the introduced species *D*_*i*_ and with increasing growth rate *r*_*i*_, while it decreases when the carrying capacity of the native population *K*_*n*_ and its competitive effect *α*_*ni*_ on the introduced population are large.

#### E.2 Speed of invasion and microbiome transfer

We interested in understanding under which circumstances microbiome transfer from natives to introduced individuals may occur before the displacement of natives. In Eq. (29) we know the speed at which the introduced population displace a native one. We can therefore calculate the time Δ*t* needed for the introduced species to displace natives within a certain one-dimensional patch of length Δ*x*, with invasion speed *c*^*\**^. Thus, we find that

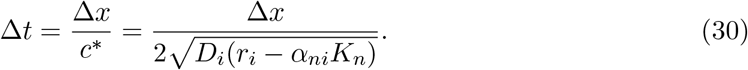

We define *t*_*d*_ as the minimal average time needed for the first microbiome transfer event to occur, which depends on the expected value of the microbiome transfer rate between native and introduced individuals (Eq. (2)). Thus we obtain

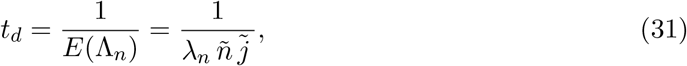

where 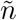 and 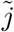are the averaged population densities of introduced and natives species at the wavefront. As long as *t*_*d*_ *<* Δ*t*, microbiome transfer from natives to the introduced population can occur before the displacement of natives. We obtain therefore a lower bound for the density-dependent microbiome transfer rate *λ*_*n*_, namely

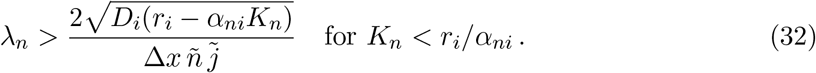

Microbiome transfer is therefore more likely to happen when the patch size Δ*x* is large, the density of natives *N* and introduced species *I*_0_ are large, and the competitive effect of natives on introduced individuals (*α*_*ni*_) is large. Microbiome transfer is less likely to occur when the patch size Δ*x* is small, and when the dispersal ability and growth rate of the introduced species are low.

**Fig. S7:**
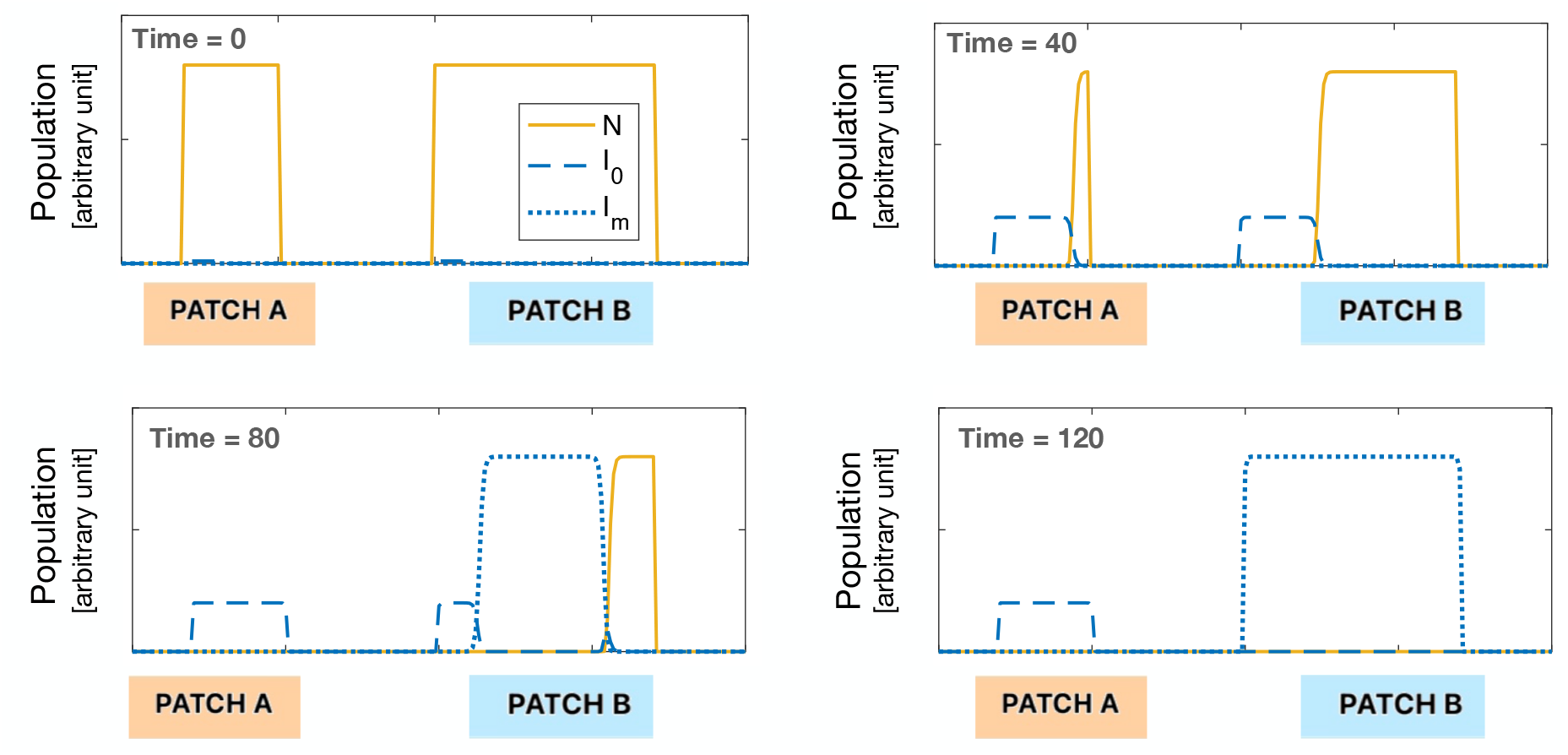
Patch A is smaller and presents a low density of natives, and the invaders displace the native population before the transfer of beneficial microbes from native to invaders can occur. In patch A, the invasive population remains therefore poorly adapted to the new environment, and in low density. Patch B presents a larger patch size and a higher density of natives, which increase the likelihood that native microbes will be transferred to the invaders before natives are driven to extinction. The acquisition of native microbes confers local adaptation to the invaders, and their population in patch B grows larger than in patch A. The findings represented in this figure are based on Eq. (5).

